# CRISPR-Cas9 induces large structural variants at on-target and off-target sites *in vivo* that segregate across generations

**DOI:** 10.1101/2021.10.05.463186

**Authors:** Ida Höijer, Anastasia Emmanouilidou, Rebecka Östlund, Robin van Schendel, Selma Bozorgpana, Lars Feuk, Ulf Gyllensten, Marcel den Hoed, Adam Ameur

## Abstract

To investigate the extent and distribution of unintended mutations induced by CRISPR-Cas9 *in vivo,* we edited the genome of fertilized zebrafish eggs and investigated DNA from >1100 larvae, juvenile and adult fish in the F0 and F1 generations. Four guide RNAs (gRNAs) were used, selected from 23 gRNAs with high on-target efficiency *in vivo* in previous functional experiments. CRISPR-Cas9 outcomes were analyzed by long-read sequencing of on-target sites and off-target sites detected *in vitro.* In founder larvae, on-target editing of the four gRNAs was 93-97% efficient, and three sites across two gRNAs were identified with *in vivo* off-target editing. Seven percent of the CRISPR-Cas9 editing outcomes correspond to structural variants (SVs), i.e., insertions and deletions ≥50 bp. The adult founder fish displayed a mosaic pattern of editing events in somatic and germ cells. The F1 generation contained high levels of genome editing, with all alleles of 46 examined F1 juvenile fish affected by on-target mutations, including four cases of SVs. In addition, 26% of the juvenile F1 fish (n=12) carried off-target mutations. These CRISPR-induced off-target mutations in F1 fish were successfully validated in pooled larvae from the same founder parents. In conclusion, we demonstrate that large SVs and off-target mutations can be introduced *in vivo* and passed through the germline to the F1 generation. The results have important consequences for the use of CRISPR-Cas9 in clinical applications, where pre-testing for off-target activity and SVs on patient material is advisable to reduce the risk of unanticipated effects with potentially large implications.

## Introduction

Genome editing using the CRISPR-Cas9 system has become an indispensable tool across many areas of biomedical research, and holds promise to revolutionize the treatment of genetic disorders^1–4^. However, the use of CRISPR-Cas9, in particular for human germline gene editing, has raised ethical questions that need careful consideration. One major aspect of attention is unintended mutations, caused by CRISPR-Cas9, at locations in the genome other than the targeted site^5, 6^. Such *off-target* mutations can have serious consequences as they might disrupt the function or regulation of non-targeted genes. In addition, larger structural changes of the genome sequence, occurring at the intended *on-target* editing site, are another cause of concern. Undesired outcomes of CRISPR-Cas9 genome editing have been the subject of many investigations. The conclusions from these studies have been somewhat conflicting, with adverse effects of CRISPR-Cas9, i.e., larger on-target SVs and off-target mutations, being reported in some cases^5–8^ but not in others^9, 10^. These discrepancies can, at least partly, be explained by differences in experimental factors such as the Cas9 concentration, delivery method, or specific properties of the cells being investigated. In other cases, limitations of the experimental setup or the genomics technologies used to interrogate the editing sites, could hinder the discovery of CRISPR-Cas9-induced events. Moreover, the adverse CRISPR-Cas9 effects may be rare and only occur in a small fraction of the edited samples. Therefore, in order to conclusively determine the effects of CRISPR-Cas9 and their long-term consequences *in vivo*, a large number of samples needs to be followed through development and over generations, using a sensitive method for genome analysis.

Validation of genome editing is often performed using short-read or Sanger sequencing. While such methods are capable of detecting small insertion and deletion events, which are the most common outcomes of CRISPR-Cas9 directed non-homologous end joining (NHEJ), they may fail to detect larger genome aberrations. Long-read sequencing does not suffer from these limitations. In a pioneering study by Kosicki et al., large deletions and complex rearrangements were shown to exist at the on-target site of genome-edited cells, through a combination of long-range PCR and long-read sequencing^11^. Following this study, Cas9-induced structural variants (SVs) have been detected *in vivo* at the on-target site^12–14^. Recently, reports have emerged describing other types of complex genome rearrangements at the on-target site, including segmental or whole chromosome deletions^15–19^ as well as chromothripsis^20^.

Large SVs and complex genome aberrations induced by CRISPR-Cas9 have so far only been systematically examined at the on-target sites. If such events also occur at off-target sites, then that would arguably be even more worrying, because large genome aberrations in chromosomal regions or genes not intended or monitored for editing could lead to unpredictable functional consequences. To examine whether large SVs at off-target sites are cause for concern, their genomic locations first need to be established. The off-target locations can be predicted by computational tools^21–25^, but a more reliable approach is to experimentally determine the Cas9 off-target activity *in vitro* using a sequencing assay^26–31^. For this purpose, we recently developed Nano-OTS, a long-read sequencing assay based on nanopore sequencing^32^. The Nano-OTS method does not suffer from amplification bias, and reliably identifies off-target sites, even in repetitive and complex regions of the genome.

In the present study, we aim to gain a better understanding of unintended CRISPR-Cas9 genome editing outcomes at on- and off-target sites *in vivo*, and in particular SVs that may escape detection by short-read or Sanger sequencing. To accomplish this, we obtained DNA from a large number of CRISPR-Cas9-edited zebrafish and their offspring, and examined the on-target and off-target sites using long-read sequencing. Though our study was performed in zebrafish, we expect that the findings can be extrapolated also to other vertebrate animals, including humans, provided that the editing experiments are performed under similar conditions. Finally, we propose a new strategy for detection and validation of CRISPR-Cas9 genome editing outcomes using long-read sequencing technology, which we believe will represent an important step towards reducing the risk of adverse effects of CRISPR-Cas9 in clinical applications.

## Results

### Detection of Cas9 off-target cleavage sites in zebrafish DNA

The aim of this study is to investigate the prevalence and distribution of different types of mutations post CRISPR-Cas9 editing at the intended target site (on-target) as well as at *in vitro-* established off-target sites. To select gRNAs for our experiments, we pre-screened 23 gRNAs with high *in vivo* on-target efficiency. These gRNAs target zebrafish *(Danio rerio)* orthologues of human genes in loci identified by genome-wide association studies (GWAS) for cardiometabolic risk factors and diseases^33–35^ and have been used to examine the role of the candidate genes in cardiometabolic disorders^36, 37^ We first used genome-wide Nano-OTS^32^ to identify off-target Cas9 cleavage activity *in vitro* for all 23 gRNAs (Supplementary Table S1). The four gRNAs with the highest number of *in vitro* detected off-target sites, and with at least one of those located within a gene, were selected for further experiments. The four selected gRNAs target early exons of *ldlra, nbeal2, sh2b3* and *ywhaqa*. Nano-OTS identified five off-target sites for the *ldlra* gRNA (three intronic), 13 for the *nbeal2* gRNA (seven intronic; two exonic), two for the *sh2b3* gRNA (one intronic) and seven for the *ywhaqa* gRNA (one intronic; six exonic) (Figure 1). The off-target sites had between two and seven mismatches with the gRNA sequence, including mismatches at the PAM site *(nbeal2* off-target 5 and *sh2b3* off-target 1). Further investigation of the off-target sites with mismatches in the PAM sequence revealed an adjacent PAM site located 1 bp downstream of the intended PAM site, with the mismatched base pair being the first nucleotide of the NGG PAM site motif.

**Figure 1.**
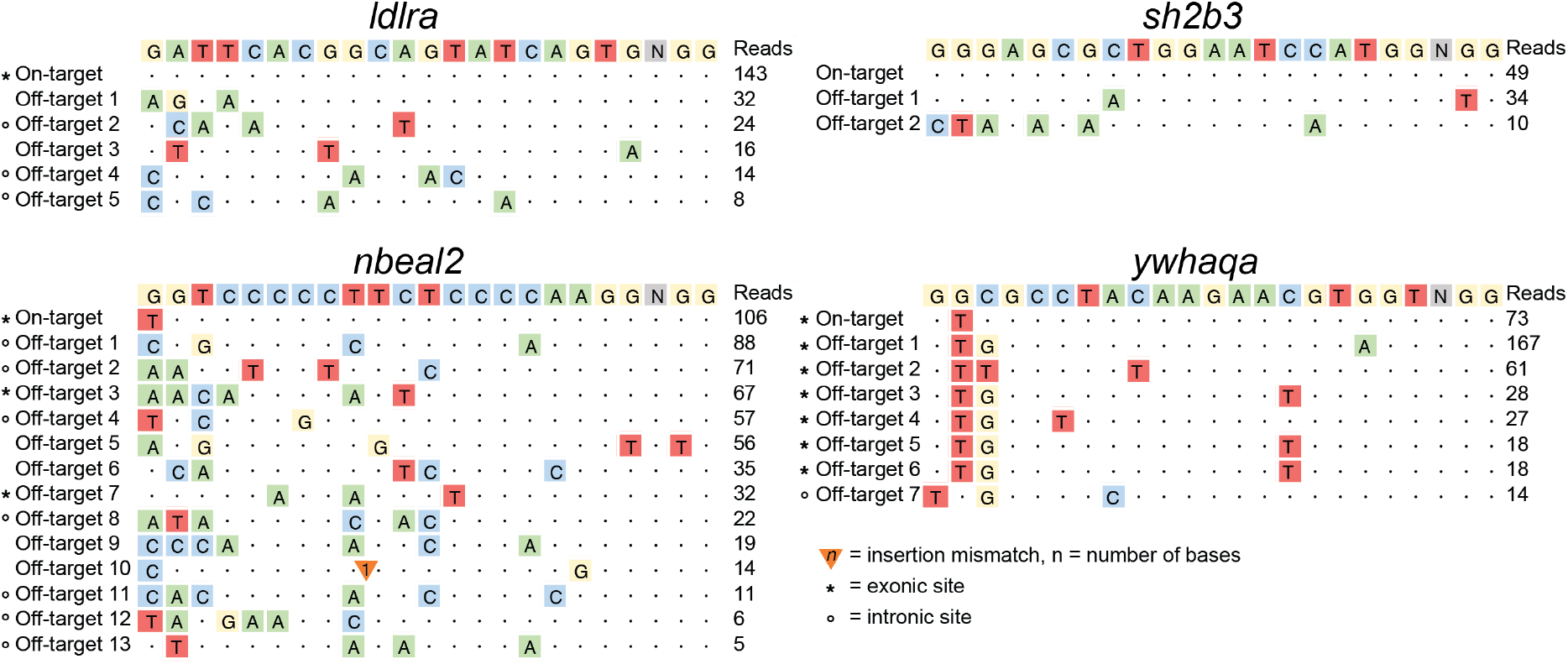
Predicted off-target sites of four guide RNAs for zebrafish genome editinsg. The diagrams show Cas9 cleavage sites detected *in vitro* by Nano-OTS for the four gRNAs *ldlra, nbeal2, sh2b3* and *ywhaqa.* The sequence at the top of each diagram displays the gRNA sequence and PAM site (NGG). The rows below show the on-target site as well as the identified off-target sites. Colored letters correspond to single-nucleotide mismatches between the target site and the GRCz11 genome. Triangles are used to mark insertion mismatches, where nucleotides need to be inserted to match the reference genome. Asterisks (*) and circles (°) mark off-target sites located within exonic and intronic regions, respectively. The column to the right shows the number of reads in the Nano-OTS analysis for each target site.

### CRISPR-Cas9 genome editing and crossing of founders

CRISPR-Cas9 editing experiments were set up as outlined in Figure 2, using the four gRNAs targeting *ldlra, nbeal2, sh2b3* and *ywhaqa.* To this end, fertilized eggs were microinjected with ribonucleoproteins (RNPs) at the single-cell stage, while uninjected eggs from the same crossing were used as controls. Microinjected RNPs typically result in >90% editing efficiency and have become a method-of-choice for genome editing in functional studies in zebrafish^38^. Samples were collected for analysis at the larval stage (5 or 10 days post-fertilization) as well as when founder fish reached adulthood (3 months). Next, we crossed randomly selected pairs of adult F0 fish to obtain an F1 generation of edited zebrafish. In the F1 generation, we collected samples at the larval stage as well as from juvenile fish (2 months). Three replicate CRISPR-Cas9 genome editing and crossing experiments were performed consecutively, using fertilized eggs from different parents of the same zebrafish line (ABs). A complete list of all examined zebrafish samples is provided in Supplementary Table S2. Since the genome editing was successful in all replicate experiments, with an on-target editing efficiency of at least 84%, all samples collected at the same developmental stage and edited with the same gRNA were jointly analyzed.

**Figure 2.**
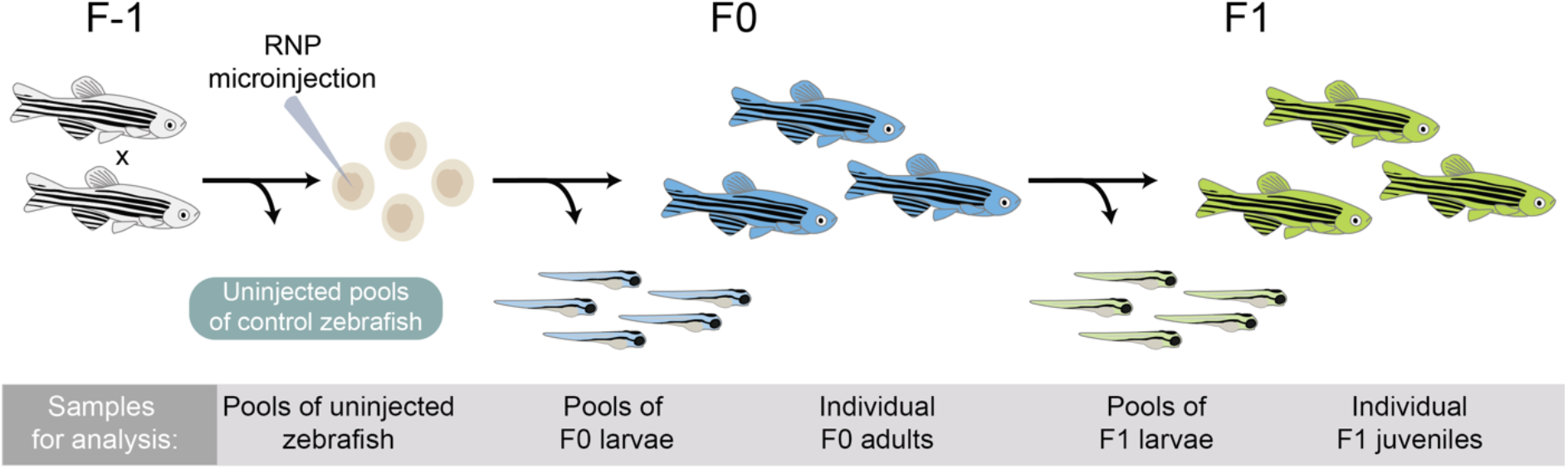
Overview of CRISPR-Cas9 genome editing in zebrafish. Genome editing was performed in fertilized eggs by microinjection of ribonucleoprotein (RNP) at the single-cell stage. The genome editing experiment results in mosaic heterozygous mutants, mosaic homozygous mutants or unaffected homozygotes. A number of F0 embryos were not injected and used as controls. F1 generation zebrafish were generated by in-crossing randomly selected pairs of adult founders. The offspring of these crossings have stable genotypes with 0, 1 or 2 mutated alleles. Samples were collected for analysis at different stages of the experiment as described in the gray box.

### CRISPR-Cas9 induces editing at on- and off-target sites

To investigate the types of Cas9-induced mutations at on- and off-target sites, we constructed large amplicons (2.6-7.7 kb) spanning the Cas9 cleavage sites in samples from edited zebrafish, as well as in uninjected controls (Supplementary Table S3). The PCR products were sequenced using the PacBio Sequel system, to obtain long and highly accurate (>QV20) reads. Detecting and quantifying genome editing outcomes from the resulting PacBio reads was performed using the software SIQ (Methods). To filter out false positives, which for example could occur due to alignment difficulties in homopolymer regions, all events detected in an uninjected control sample were removed from further analyses. On-target editing efficiencies were then calculated based on the remaining insertion or deletion mutations in the pools of founder larvae. This resulted in 92.6% on-target editing efficiency for *ldlra;* 96.7% for *nbeal2;* 92.6% for *sh2b3;* and 93.6% for *ywhaqa* (Figure 3a). In addition, we identified Cas9 activity at three off-target sites; *sh2b3* off-target 1 (editing efficiency 1.8%); *ywhaqa* off-target 1 (2.4%); and *ywhaqa* off-target 2 (6.3%) (Figure 3b).

**Figure 3.**
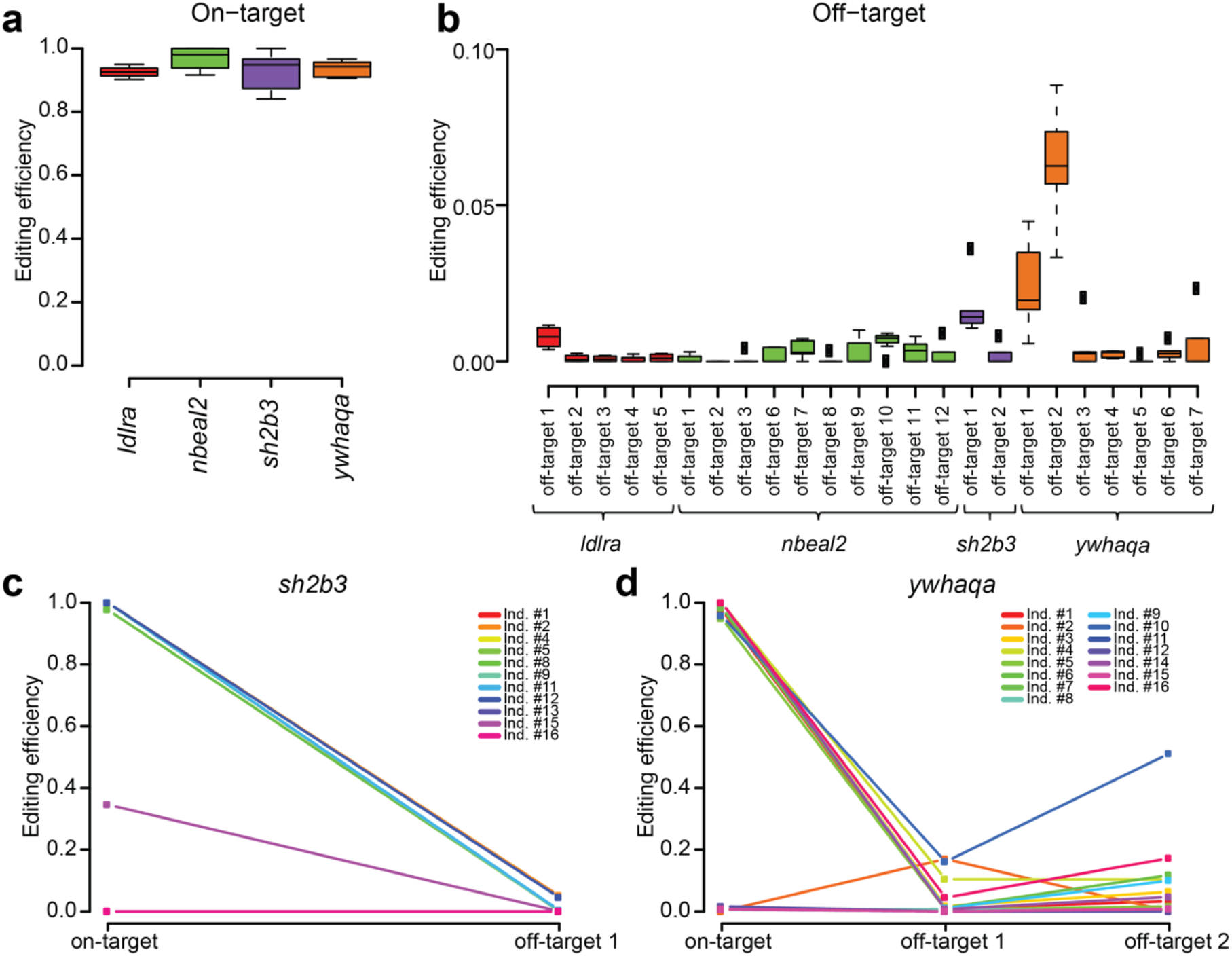
CRISPR-Cas9 editing efficiencies in pooled founder larvae and individuals. **a**) Boxplot showing on-target CRISPR-Cas9 editing efficiencies in pools of founder larvae for the four gRNAs used. **b**) Boxplot showing CRISPR-Cas9 editing efficiencies at the *in vitro-* detected off-target sites in pools of founder larvae. **c-d**) Line graphs visualizing the CRISPR-Cas9 editing efficiencies at on- and off-target sites in individual adult founders. Each colored line shows the editing efficiencies at the on- and off-target site(s) in one individual.

### Founder fish are highly mosaic in somatic and germ cells

We next examined the on-target and the three *in vivo*-confirmed off-target sites in 26 adult founders at three months of age, edited either for *sh2b3* (n=11) or *ywhaqa* (n=15) at the single cell stage. 18 of 26 F0 fish (69.2%) showed on-target editing (Figure 3c-d). Many distinct insertion and deletion events, as well as more complex combinations of insertions and deletions, were observed in any single individual, consistent with mosaicism of genome editing outcomes at the on-target site (Figure 4). To facilitate the downstream analyses, every event that consisted of a combined insertion and deletion was counted either as an insertion or a deletion, depending on which of the two sub-events that involved the highest number of nucleotides. By further examining CRISPR-Cas9-induced mutations in individual F1 juvenile fish and pooled larvae from the same founder parents, we noticed that up to six unique alleles were passed on from a single F0 breeding pair (Supplementary Tables S4-S9). Since at most four alleles are expected at any given locus, this observation is consistent with mosaicism in the founders’ germ cells. Six founder fish (23%) displayed off-target genome editing in at least 10% of DNA molecules, with the highest proportion (50.4%) observed in individual #10 at *ywhaqa* off-target 2 (Figure 3d).

**Figure 4.**
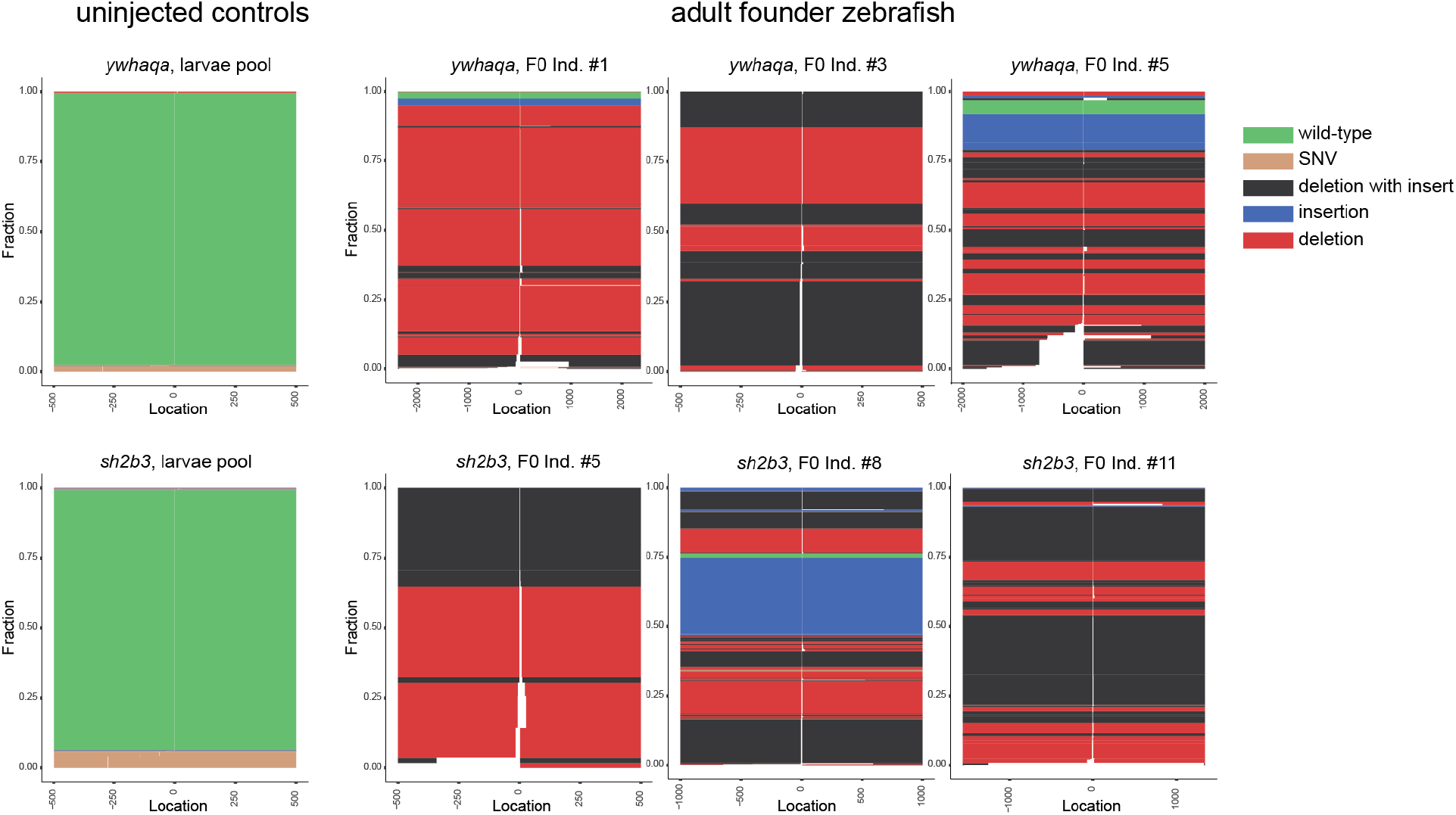
Somatic mosaicism of CRISPR-Cas9 editing events in founder fish. Schematic view of editing events at the on-target sites for *ywhaqa* (top) and *sh2b3* (bottom), produced by the SIQ software. To the left are two control samples from uninjected zebrafish, where only wild-type alleles and sequences containing SNVs are reported. To the right are results from three individual F0 founder fish edited for *ywhaqa* or *sh2b3.* In the CRISPR-Cas9 edited F0 individuals, a large number of distinct insertions, deletions, as well as combinations of insertions and deletions are reported. This is consistent with somatic mosaicism of CRISPR-Cas9 editing events in the founder fish.

### Large structural variants at on- and off-target sites

To determine the size distribution of genome editing events induced by CRISPR-Cas9 *in vivo,* we focused on pools of founder larvae. A total of 595 larvae were analyzed in 20 pools, thereby giving a comprehensive view of the different alleles introduced by CRISPR-Cas9 at an early developmental stage. The on-target events ranged from 4.8 kb deletions to 1.4 kb insertions (Figure 5a-c). Although the majority of events were small, 7% represent SVs of ≥50 bases. A similar size distribution was seen at the three off-target sites for *sh2b3* and *ywhaqa*, even though the lower degree of off-target editing results in a diagram with fewer data points (Figure 5d-f). Large SVs were detected not only in founder larvae, but also in founder adults. Strikingly, one 903 bp deletion at an off-target site completely removes an exon of a gene that was not intended to be targeted in the experiment (i.e. *ywhaqb,* Figure 5g).

**Figure 5.**
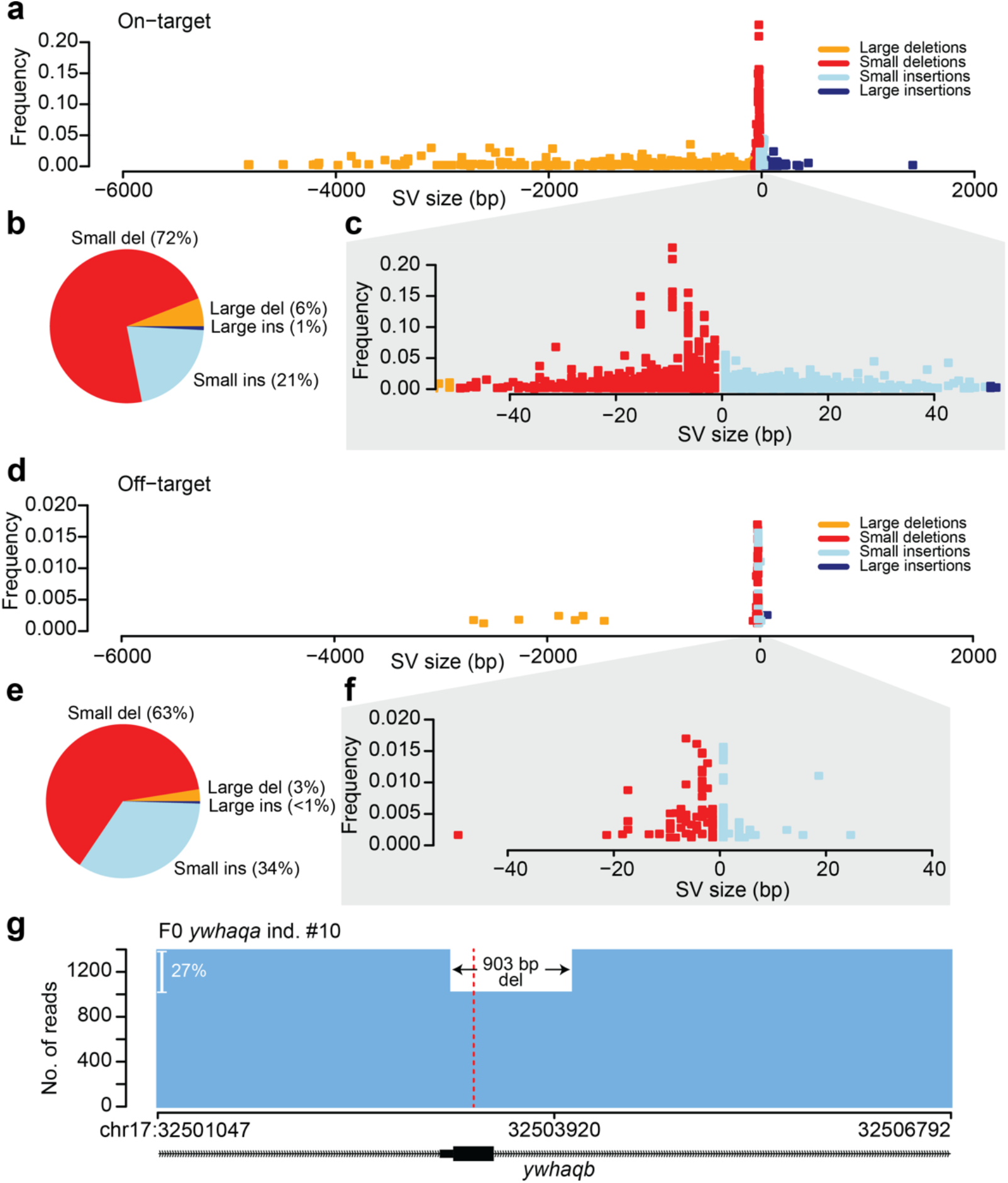
Size distribution of CRISPR-Cas9-induced mutations. **a**) A dot plot showing the size distribution of Cas9-induced on-target mutation events (large and small insertions and deletions) in pools of founder larvae. Each point in the graph visualizes a specific event in a sample, where the x-axis displays the size of the variant (negative values for deletions and positive values for insertions), and the y-axis displays the frequency of the variant in the individual sample. **b)** Fractions of small and large insertions and deletions at on-target sites in pools of founder larvae. **c)** Zoomed-in view of the plot in a), only visualizing small insertions and deletions. **d)** Size distribution of Cas9-induced off-target mutations *(sh2b3* off-target 1, *ywhaqa* off-target 1 and 2) in pools of founder larvae. **e)** Fractions of small and large insertions and deletions at the three off-target sites in pools of founder larvae. **f)** Zoomed-in view of the plot in d), only visualizing small insertions and deletions. **g)** An example of an adult founder fish with a 903 bp deletion at *ywhaqa’s* off-target 2 that spans an entire exon of *ywhaqb.* The coverage plot shows the number of reads with the 903 bp deletion and the number of reads that lack the deletion (i.e., unmodified and other Cas9-induced variants). The Cas9 cleavage site is indicated by the red line.

### Off-target mutations and SVs can be passed to the F1 generation

We next compared the frequency of genome editing events over developmental stages and generations. The proportion of edited alleles was higher in the F1 generation, where all 46 juvenile individuals were completely edited, as compared to the F0 generation where eight fish showed little or no editing (Figure 6a). Structural variants of ≥50bp are also more abundant in the F1 generation (Figure 6b). Four of the 46 juvenile F1 individuals (9%) were hetero-or homozygous for an on-target SV. Editing events at the three off-target sites showed a similar pattern, with 12 of 46 F1 fish (26%) displaying editing in at least 20% of the reads i.e., representing hetero- and homozygous individuals, as compared with adult founders where six of 26 fish (23%) displayed off-target editing at a 10% level (Figure 6c). No SVs ≥50bp were detected at off-target sites in the F1 generation (Figure 6d).

**Figure 6.**
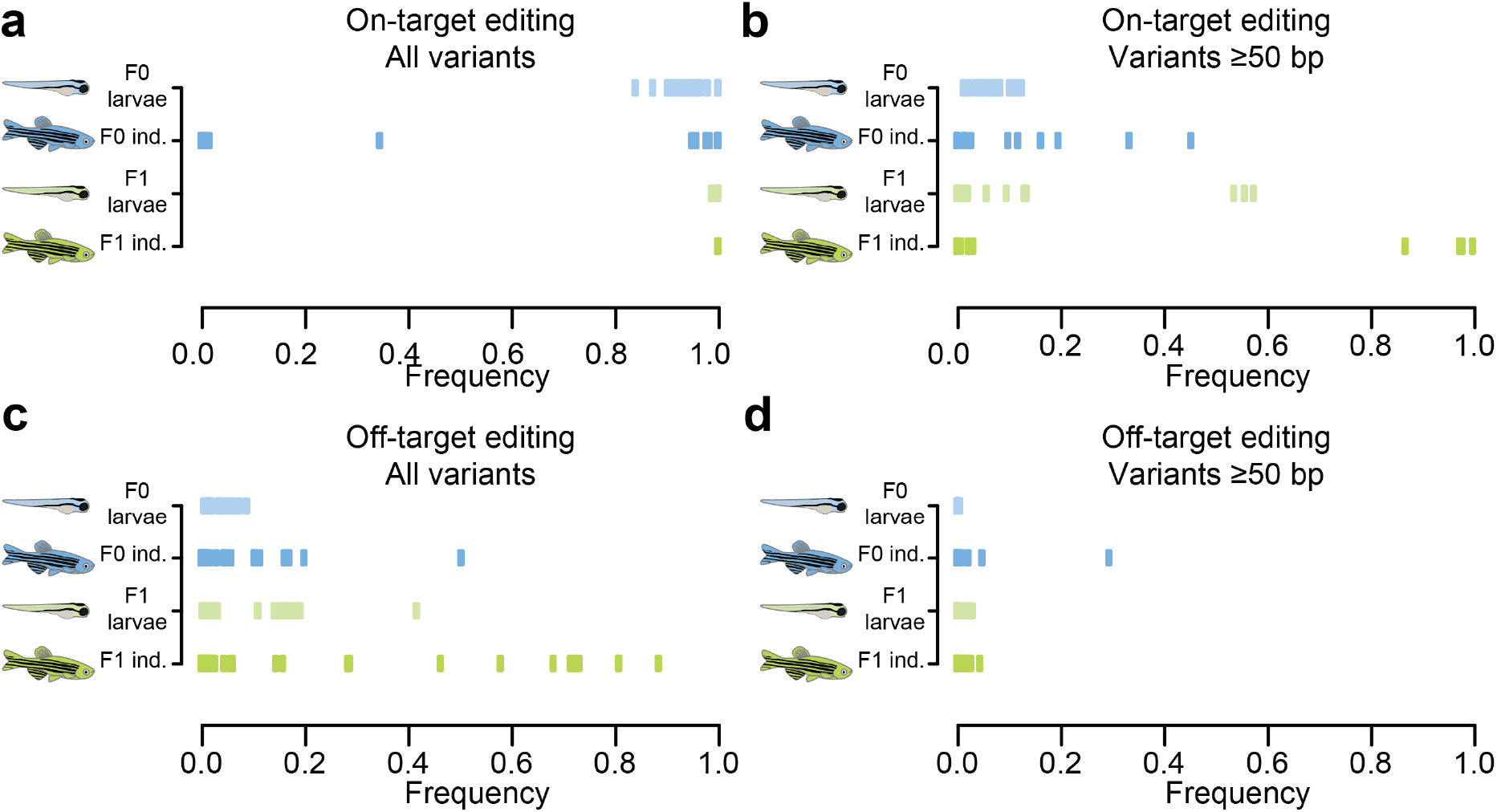
Summary of Cas9-induced variants at on- and off-target sites. Summarizing plots displaying the frequencies of Cas9-induced variants in all analyzed samples for the F0 and F1 generations (a-d). One point in the plot represents one sample (i.e., a pool of larvae or an individual juvenile/adult fish). **a)** The total frequencies of Cas9-induced variants at on-target sites. **b)** The frequencies of SVs induced by Cas9 at on-target sites. **c)** The total frequencies of Cas9-induced variants at off-target sites *(sh2b3* off-target 1, *ywhaqa* off-targets 1 and 2). **d)** The frequencies of SVs induced by Cas9 at off-target sites.

### Validation of unintended CRISPR-Cas9 editing in F1 fish

As mentioned above, we identified four juvenile F1 fish with a ≥50 bp SV at an on-target site, and 12 F1 individuals with smaller off-target mutations. Our experimental setup enabled us to search for the same events in pools of F1 larvae from the same parents, as well as in other F1 siblings. This way, we were able to verify that all unintended on-target and off-target mutations indeed exist in related larvae and juvenile fish (Supplementary Table S10). Figure 7a-b shows the results for two F1 individuals with large on-target SVs. A 1053 bp deletion, removing a big fraction of the targeted exon of *sh2b3* was found in a juvenile F1 fish as well as in pooled F1 larvae (Figure 7a,c). Furthermore, a large 292 bp insertion in the targeted exon of *ywhaqa,* observed in three juvenile F1 fish, was also detected in pooled F1 larvae (Figure 7b,d). Unexpectedly, for 7 of the 46 F1 individuals (15%), >98% of the reads support only one specific editing event (Supplementary Table S11). These fish could be homozygous for the edited locus^39^, carry a large deletion on the other chromosome, or, alternatively, a different allele exists that was not detected due to allelic dropout. While we cannot with certainty provide the reason for this frequently observed homozygosity, our results confirm that large SVs and off-target mutations are not only observed in founder fish, but also in the F1 generation.

**Figure 7.**
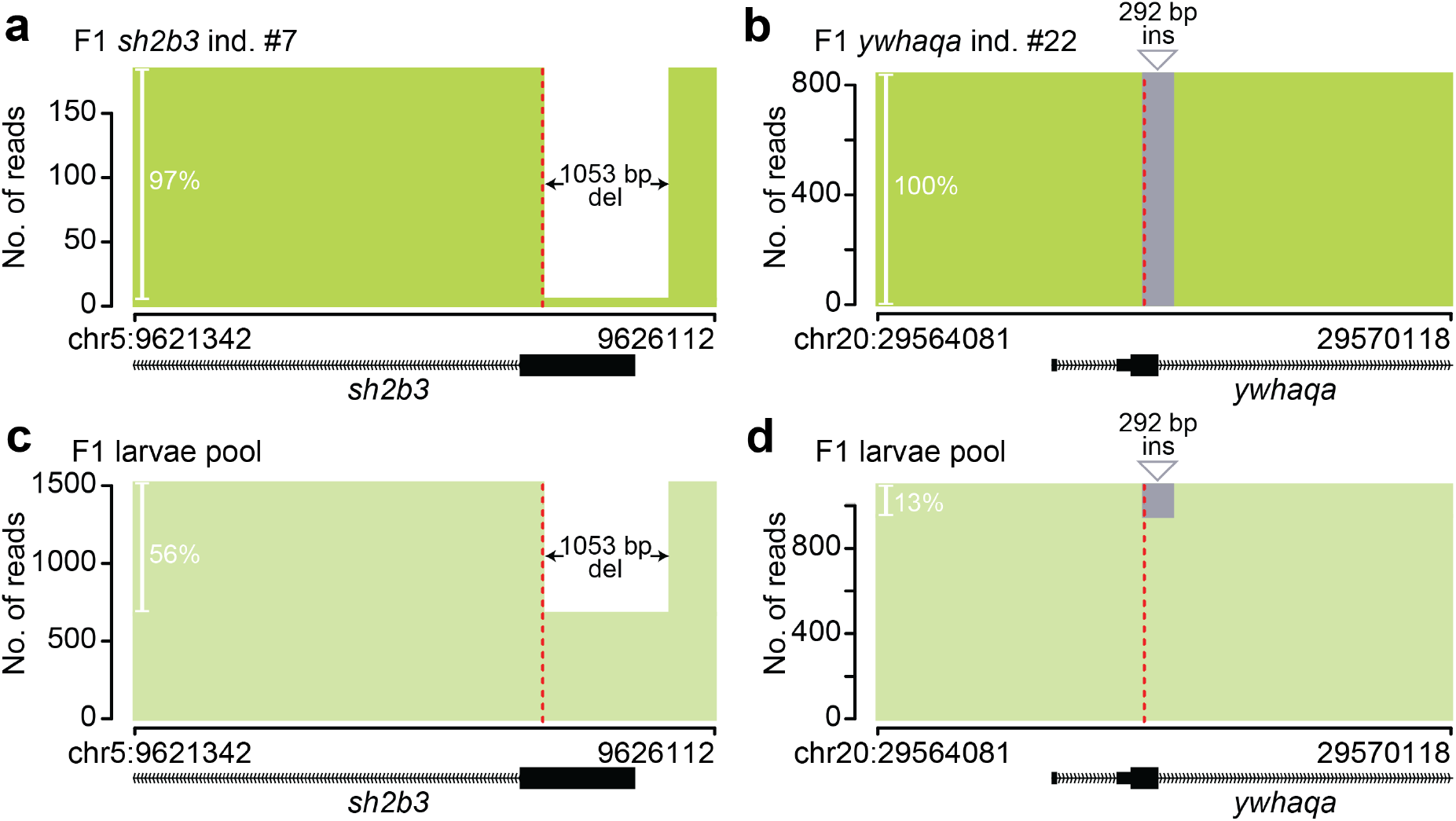
Large structural variants in individual F1 zebrafish. **a-b)** Examples of large SVs at on-target sites in individual juvenile F1 zebrafish, a 1053 bp deletion in *sh2b3* (a) and a 292 bp insertion in *ywhaqa* (b). **c-d)** The same variants are observed in pools of F1 larvae from the same F0 pair as the individuals in a) and b). The plots show the number of reads with the large SV and the reads without the variant (i.e., unmodified and other Cas9-induced variants). The Cas9 cleavage sites are indicated by red lines.

## Discussion

In this study, we used long-read sequencing to examine on- and off-target genome editing outcomes, across multiple stages of development and across generations. Genome editing was accomplished using standard routines for germline editing at the single-cell stage, using gRNAs that have recently been used for functional studies of cardiometabolic diseases in zebrafish model systems^36, 37^. This revealed insertions and deletions of sizes up to several kilobases, at on- and off-target sites, with a high degree of individual-level variation in genome editing outcomes both in the F0 and F1 generations. A major advantage of our experimental setup is that editing events detected in the F1 generation could be directly verified in siblings from the same parents. In this way we were able to validate all off-target mutations and larger SVs in the F1 generations. At the same time, we confirmed the absence of events in uninjected controls and in F1 fish from different founders. We can therefore conclude that no false positives were introduced by DNA amplification, long-range sequencing or the downstream analysis.

In the founder larvae, about 7% of the editing outcomes correspond to insertions or deletions of ≥50 bp. This number gives a first estimate of the abundance of SVs during early cell division. At the on-target site, several large SV events can also be observed in adult founders, whereafter they in some instances segregate to the next generation. Four of the 46 juvenile F1 fish we examined carried a large deletion or a large insertion at the on-target site. We also observed a 903 bp deletion at an off-target site that removes an exon of *ywhaqb* in one F0 individual. Unwanted large SVs in coding regions are problematic, but even more so when they occur at off-target sites, where they likely remain undetected.

Our data also point to several unexpected features of the CRISPR-Cas9 system that warrant further investigation. Firstly, we find that the germ cells of founder fish are mosaic. This could help us understand how CRISPR-Cas9 events are inherited to the next generation. Moreover, 15% of the juvenile F1 fish seem to be homozygous for one specific CRISPR-Cas9 editing event, while the remaining ones are compound heterozygous. For the homozygous F1 fish, either one of the alleles failed to be amplified by the long-range PCR, or alternatively there exists two distinct alleles with identical editing events. Through additional experiments and analyses it might be possible to determine if there are other, possibly even larger events induced by CRISPR-Cas9 that result in failed amplification of the affected allele; if there is loss-of-heterozygosity within the region; or if the homozygosity is driven by non-random DNA repair resulting in the same mutations across multiple founders^39^. In any case, based on our results, more experiments are required to further improve our understanding of the multigenerational genomic consequences of CRISPR-Cas9 editing.

Even though this work is based on DNA from a large number of individuals, and using modern genomics technologies, many factors and parameters in the experiment could have influenced the results. For example, we microinjected RNPs to get the CRISPR-Cas9 machinery into the cells. Different results might have been obtained if we had used other methods for transfection. Additional factors such as differences in DNA repair system between cell types, or the total concentration of Cas9 within the cells, could also influence the editing efficiency and outcomes at the on- and off-target sites. For these reasons, it would be desirable to perform similar experiments also in other sample types and organisms. A further limitation of our study is that even long-range PCR is unable to capture large genome aberrations and complex genomic rearrangements, such as chromothripsis^20^ and whole chromosome deletions^19^. Detecting such events requires an additional analysis of the edited samples that either studies the whole genome, or an alternative method for enrichment and long-read sequencing^40–42^. However, since we require viable outcome for the zebrafish, it is unlikely that large and complex genome alterations will be heritable and present in the analyzed samples.

Our finding that CRISPR-Cas9 can induce large SVs at on- and off-target sites *in vivo* does not mean we should stop using this powerful tool. For genetic screens in cellular systems or for functional experiments in model organisms, the impact of these large SV events will be relatively modest, since only a limited number of individuals or samples are likely to be affected. However, for clinical applications, such as genome editing in monogenetic disorders, it is critical to identify potentially serious adverse effects caused by *a priori* unexpected genome editing in the cells of interest. Lastly, when it comes to manipulation of human embryos, our study adds yet more arguments for caution, due to the unintended mutations that can have consequences for the individual and, in some cases, future generations.

In conclusion, by applying new genomics tools and carefully designed experiments, we can learn more about the consequences of CRISPR-Cas9 editing *in vivo,* while at the same time developing improved strategies to validate edited cells. Based on our findings, we propose the following three-step approach to verify CRISPR-Cas9 genome editing outcomes for clinical applications: i) employ an *in vitro* method, such as Nano-OTS, to detect where Cas9 cleavage sites are located in DNA from the individual and – if possible – cell type of interest, ii) perform long-read re-sequencing of the on-target and predicted off-target sites, to determine the genotypes in each individual sample, and iii) ensure that there is no allelic dropout in the region, through identification of nearby polymorphic variants or by further experimental verification.

This three-step approach enables detection of unintended SVs that are missed with short-read assays, and is therefore an important step towards a safer use of CRISPR-Cas9 for therapeutic purposes.

## Acknowledgements

Sequencing was performed by the SciLifeLab National Genomics Infrastructure (NGI) in Uppsala, Sweden. Computations were performed on resources provided by SNIC through Uppsala Multidisciplinary Center for Advanced Computational Science (UPPMAX) under Project SNIC 2021/23-81. We would like to thank Mai-Britt Mosbech, Uppsala University, for advice and recommendations for high molecular weight DNA extractions in zebrafish. This study was supported by grants from Swedish Cancer Foundation and Swedish Research Council (U.G.). M.d.H is supported by grants from the Swedish Heart-Lung Foundation (20200781, 20200602), the Kjell and Märta Beijer Foundation, and the Swedish Research Council (2019-01417).

## Authors’ contributions

IH, LF, UG, MdH and AA conceived the study. IH and AA drafted the manuscript. IH, AE and RÖ performed the experiments. IH, RvS, SB and AA performed the analyses. All authors read and approved the final manuscript.

## Data availability

All sequencing data will be made available from the NCBI Sequence Read Archive (SRA).

## Methods

### Zebrafish handling and CRISPR-Cas9 genome editing

All zebrafish experiments and husbandry were conducted in accordance with Swedish and European regulations, and have been approved by the Uppsala University Ethical Committee for Animal Research (Dnr 5.8.18-13680/2020). The genes of interest were targeted using one gRNA per orthologue that had an anticipated efficiency >90%. RNA duplexes of the chemically synthesized Alt-R^®^ crRNA (IDT) and Alt-R^®^ tracrRNA (IDT) were complexed with Alt-R^®^ S.p. Cas9 nuclease, v.3 (IDT) to form “duplex guide RNPs” (dgRNPs), as decribed by Hoshijima K et al.^38^. The dgRNPs were then injected into fertilized zebrafish eggs at the 1-cell stage. Uninjected embryos were kept and used as controls. The injected founder embryos were raised to adulthood at which time random mating pairs were in-crossed. Pools of 25-30 five- or ten-day-old F0 larvae, individual F0 adult fish, pools of 30 five-day-old F1 larvae and fin clips individual F1 fish were collected throughout the experiment for downstream analyses. The complete sample collection is described in Supplementary Table S2. Adult zebrafish were sacrificed by prolonged exposure to tricaine, followed by snap freezing in liquid nitrogen to ensure DNA integrity.

### Extraction of genomic DNA from zebrafish

All samples were extracted using the MagAttract HMW DNA Kit (Qiagen) and the “Manual Purification of High-Molecular-Weight Genomic DNA from Fresh or Frozen Tissue” protocol according to manufacturer’s instructions. A tissue homogenization step using a pestle was added to the protocol prior to the lysis step. DNA integrity of the extracted samples was assessed using the Femto Pulse system (Agilent Technologies) using the Genomic DNA 165 kb kit.

### Detection of off-target sites using Nano-OTS

Genomic DNA was sheared to 20 kb fragments using the Megaruptor 2 (Diagenode) and size selected with a 10 kb cut-off using the BluePippin system (Sage Science). 4 μg of sheared and size-selected DNA was then used for Nano-OTS library preparation described by Höijer et al. ^32^. To increase coverage two libraries were prepared and sequenced on one R9.4.1 flow cell each. Guppy v4.0 was used for base calling.

### Alignment of reads and detection of off-target sites

The reads from Nano-OTS were aligned to the GRCz11 reference genome using minimap2^43^, after which the Cas9 cleavage sites were predicted using v1.8 of the Insider software (https://github.com/UppsalaGenomeCenter/InSiDeR). For each predicted Cas9 cleavage site, the corresponding sequence from GRCz11 was extracted in a +-40 bp window surrounding the Cas9 cleavage site. All sequences containing gaps (N’s) were filtered out since we were only interested in detecting gRNA binding events in high-quality regions of the zebrafish genome. For the remaining sequences, we globally aligned against all gRNA sequences using v6.6.0 of EMBOSS-Needle with default settings^44^. Only sequences with containing an alignment score of >55 to a certain gRNA were considered positive binding sites.

### Amplicon construction and sequencing

Primers were designed for all on-target and off-target sites predicted by Nano-OTS for the four gRNAs. Primers for three off-target sites for the *nbeal2* gRNA were excluded due to PCR optimization difficulties or because of issues with the GRCz11 zebrafish reference genome. The amplicons range from 2.6 kb to 7.7 kb in size. Primer sequences, expected amplicon sizes and primer coordinates can be found in Supplementary Table S3. Long-range PCRs were performed using the PrimeStar GLX Polymerase (Takara Bio) according to manufacturer’s instructions, using either the standard or 2-step PCR cycling protocol. 0.2 μg/μl bovine serum albumin (BSA) was added to the PCR reactions for improved performance. PCRs were performed using 30 ng of genomic DNA from pooled or individual zebrafish DNA extractions. Amplicons originating from different primer pairs were pooled in an equimolar fashion. Amplicon pools were barcoded and sequenced on PacBio’s Sequel system using the SMRTbell^®^ Express Template Prep Kit 2.0 and the PacBio Barcoded Overhang Adapter Kit 8A and 8B for SMRTbell construction, and 3.0 sequencing and binding chemistry using a 10-hour movie time.

### Analysis of on- and off target mutations in long amplicon data

HiFi reads were created for the SMRT long-amplicon data, after which alignment was performed to GRCz11 using minimap2^43^. On- and off-target editing efficiencies were calculated as the fraction of reads containing insertions and deletions at the Cas9 digestion site. To ensure that indel variation at the Cas9-cleavage sites target sites is caused by CRISPR-Cas9 genome editing, and by genetic variation in the zebrafish, systematic errors introduced in the sequencing, or alignment artefacts, we removed all sites having a frequency of at least 0.5% indel mutations in the F0wt pool. HiFi reads were created for the SMRT long-amplicon data, after which alignment was performed to GRCz11 using minimap2^43^. After alignment, all reads covering a specific on-target or off-target site were analyzed using the software SIQ (manuscript in preparation). SIQ performs a detailed analysis of all reads covering a specific target and reports the identified editing events along with their frequencies. To remove potential false positive events reported by SIQ in the edited samples, all events that were also detected in the control samples were flagged and considered as unedited. Custom R scripts were used to visualize the SIQ results. In cases where an SV event simultaneously contain an insertion and a deletion, the event was visualized either as an insertion or as a deletion depending on which part of the SV had the largest size.

