## Supplementary Material for "CRISPR-Cas9 induces large structural variants at on-target and off-target sites *in vivo* that segregate across generations"

### Supplementary Tables

**Table S1.** Cas9 cleavage sites *in vitro* for 23 gRNAs, detected by the Nano-OTS method. Coordinates and sequences are obtained from the danRer11 zebrafish reference genome.

| chr | position | gRNAname | gDNAsequence | alignSimilarity | alignScore | snvBases | indelBases | peakHeight |
|---|---|---|---|---|---|---|---|---|
| chr17 | 30718007 | apoba | TGGGAGGGCTCTATCTTAGGGGG | 21/80 (26.2%) | 99 | 1 | 0 | 192 |
| chr14 | 40455513 | apoba | CCGCCTATGATAGAGCCCTAATT | 17/80 (21.2%) | 63 | 5 | 0 | 111 |
| chr22 | 36284098 | apoba | GGGGAAGGCAGTATCTTAGGGGG | 19/80 (23.8%) | 81 | 3 | 0 | 85 |
| chr20 | 31273933 | apobb1 | GCAGCTGACAAAGTACCAAGGGG | 21/80 (26.2%) | 99 | 1 | 0 | 65 |
| chr22 | 4914510 | apobb1 | CCTCTTGGTTCTTTGTACAGCTCT | 20/80 (25.0%) | 70 | 2 | 1 | 23 |
| chr20 | 53444334 | apobb2 | CCATCCTCTTGCTCAGCTAATCA | 21/80 (26.2%) | 99 | 1 | 0 | 87 |
| chr2 | 47454801 | apobb2 | CCCTCCTCTTGCTCTGCTACTCT | 19/80 (23.8%) | 81 | 3 | 0 | 24 |
| chr16 | 23985170 | apoc2 | CCTTGCACTGGGTATGTACACAC | 21/80 (26.2%) | 99 | 1 | 0 | 122 |
| chr19 | 10856069 | apoea | CCCCTTACGCGCAGGAGAGAGCC | 22/80 (27.5%) | 108 | 0 | 0 | 116 |
| chr16 | 23961550 | apoeb | CCTGTTCCAGGCTGATGCCCCTC | 22/80 (27.5%) | 108 | 0 | 0 | 209 |
| chr17 | 5878029 | gckr | CCATCCCAGATAATACACATGCG | 21/80 (26.2%) | 99 | 1 | 0 | 411 |
| chr23 | 30916318 | gckr | TGTTTGTGTATTATCTGGAAAGG | 18/80 (22.5%) | 72 | 4 | 0 | 132 |
| chr1 | 21357140 | gckr | CCTTCTCAGATGATGCACATGCA | 18/80 (22.5%) | 72 | 4 | 0 | 60 |
| chr21 | 23319983 | gckr | CCAACCCAGATAATACACCACTT | 16/80 (20.0%) | 54 | 6 | 0 | 31 |
| chr3 | 21773503 | gckr | CCGTCCCAGATGATATACAACAT | 16/80 (20.0%) | 54 | 6 | 0 | 31 |
| chr6 | 44274257 | gckr | TCTGTGTGTATTATCTGGGGACG | 16/80 (20.0%) | 54 | 6 | 0 | 19 |
| chr19 | 11525479 | gckr | GACATGTATTTTATTT------- | 12/87 (13.8%) | 44 | 4 | -1 | 10 |
| chr21 | 16260203 | gckr | ACACCCCAGACAGCACACATACG | 15/80 (18.8%) | 45 | 7 | 0 | 10 |
| chr11 | 32265767 | gckr | CCAACCCAGATAATACACCATAA | 16/80 (20.0%) | 54 | 6 | 0 | 8 |
| chr4 | 32765162 | gckr | ACTGCCCAGACAGCACACATACG | 15/80 (18.8%) | 45 | 7 | 0 | 8 |
| chr5 | 671631 | gckr | CACTCCCAGAATGTACACATGCT | 17/80 (21.2%) | 63 | 5 | 0 | 5 |
| chr14 | 48008478 | gckr | ------CGGAGAAAACCCATGCG | 12/86 (14.0%) | 40 | 5 | -1 | 4 |
| chr18 | 1143644 | hcn4 | CCGCGGCTCCAGGAGGGCTCTCC | 22/80 (27.5%) | 108 | 0 | 0 | 31 |
| chr25 | 29273090 | hcn4l | CCACGTTAAACACTATCCAGGCG | 21/80 (26.2%) | 99 | 1 | 0 | 19 |
| chr11 | 34856421 | hcn4l | CCGAGTTAAACACTATCCCCAAC | 17/80 (21.2%) | 63 | 5 | 0 | 9 |
| chr2 | 37570139 | insra | GGGACAGAGGCCAGCACAAGGGG | 22/80 (27.5%) | 108 | 0 | 0 | 147 |
| chr4 | 54914411 | insra | GGGGGAGGGGGCAGCTCCA---- | 13/84 (15.5%) | 41 | 6 | -1 | 93 |
| chr4 | 8537670 | insra | CCGCTTGTGTTGGCCTCTGACTC | 19/80 (23.8%) | 81 | 3 | 0 | 88 |
| chr23 | 14699329 | insra | TGAACCGCTGCCTGCACAAGCTG | 15/80 (18.8%) | 45 | 7 | 0 | 10 |
| chr9 | 8529080 | insra | GGAACACAGGCGGACACAAGTGC | 16/80 (20.0%) | 54 | 6 | 0 | 8 |
| chr22 | 11004684 | insrb | CCGCTCCCCGCGTGATGTTGGTC | 22/80 (27.5%) | 108 | 0 | 0 | 121 |
| chr14 | 15606594 | insrb | GAGCCCCATCGGGGGGGGGGGGG | 15/80 (18.8%) | 45 | 7 | 0 | 85 |
| chr10 | 15311488 | kcnv2a | CCTTTGGGCATACGCGTTTCCTC | 22/80 (27.5%) | 108 | 0 | 0 | 19 |
| chr3 | 19304777 | ldlra | GATTCACGGCAGTATCAGTGTGG | 22/80 (27.5%) | 108 | 0 | 0 | 143 |
| chr23 | 8189813 | ldlra | CCCCACTGAAACTGCCGTGTACT | 18/80 (22.5%) | 72 | 4 | 0 | 32 |
| chr16 | 9332405 | ldlra | GCATAACGGCTGTATCAGTGTGG | 18/80 (22.5%) | 72 | 4 | 0 | 24 |

| | | | | | | | | |
|---|---|---|---|---|---|---|---|---|
| chr10 | 36351898 | ldlra | CCTTACTGATACTGCAGTGAAAC | 19/80 (23.8%) | 81 | 3 | 0 | 16 |
| chr19 | 25977945 | ldlra | CCTCACTGATGTTGTCGTGAATG | 18/80 (22.5%) | 72 | 4 | 0 | 14 |
| chr24 | 7925197 | ldlra | CACTCACAGCAGTAACAGTGTGG | 18/80 (22.5%) | 72 | 4 | 0 | 8 |
| chr7 | 30398243 | lipca | CCTAGCTATGTCTGGGAGAACCC | 22/80 (27.5%) | 108 | 0 | 0 | 64 |
| chr15 | 25555212 | mmp20a | CCACATCTTGCCTCTTTCATCCC | 22/80 (27.5%) | 108 | 0 | 0 | 35 |
| chr16 | 16478569 | nbeal2 | CCTCCTTGGGGAGAAGGGGGACA | 21/80 (26.2%) | 99 | 1 | 0 | 106 |
| chr12 | 24373304 | nbeal2 | CGGCCCCCCTCTCCCAAAGGGGG | 18/80 (22.5%) | 72 | 4 | 0 | 88 |
| chr8 | 47074469 | nbeal2 | CCACCTTGGGGGGAAAGGAGATT | 17/80 (21.2%) | 63 | 5 | 0 | 71 |
| chr14 | 16385238 | nbeal2 | CCCCCTTGGGGAAATGGGGTGTT | 16/80 (20.0%) | 54 | 6 | 0 | 67 |
| chr19 | 44806008 | nbeal2 | CCACCTTGGGGAGAAGCGGGGCA | 19/80 (23.8%) | 81 | 3 | 0 | 57 |
| chr24 | 13828273 | nbeal2 | CACACTTGGGGAGCAGGGGGCCT | 17/80 (21.2%) | 63 | 5 | 0 | 56 |
| chr18 | 14204547 | nbeal2 | GCACCCCCTTTCCCCCCAGGCGG | 17/80 (21.2%) | 63 | 5 | 0 | 35 |
| chr10 | 1890600 | nbeal2 | GGTCCACCATCTTCCCAAGGAGG | 19/80 (23.8%) | 81 | 3 | 0 | 32 |
| chr3 | 35685641 | nbeal2 | ATACCCCCCTACCCCCAAGGGGG | 16/80 (20.0%) | 54 | 6 | 0 | 22 |
| chr20 | 24396692 | nbeal2 | CCCCCTTTGGGGGATGGGGTGGG | 15/80 (18.8%) | 45 | 7 | 0 | 19 |
| chr6 | 23360436 | nbeal2 | CGTCCCCCTGTCTCCCCAGGGCGG | 20/80 (25.0%) | 70 | 2 | 1 | 14 |
| chr14 | 11142437 | nbeal2 | CCACCTGGGGGGGATGGGGGGAG | 16/80 (20.0%) | 54 | 6 | 0 | 11 |
| chr18 | 30359856 | nbeal2 | CCCCCTTGGGGAGAGGGTTCATA | 16/80 (20.0%) | 54 | 6 | 0 | 6 |
| chr5 | 13924978 | nbeal2 | CCACCTTTGGGATATGGGGGAAC | 18/80 (22.5%) | 72 | 4 | 0 | 5 |
| chr7 | 53879247 | neo1a | GGAGCCGTCGGATACACTAGCGG | 22/80 (27.5%) | 108 | 0 | 0 | 70 |
| chr2 | 33150097 | neo1a | CCTTTAGTGTATCAGGCGGCTCT | 18/80 (22.5%) | 72 | 4 | 0 | 6 |
| chr25 | 2899718 | neo1b | GGACAGAGATGCTCGGCCTGTGG | 22/80 (27.5%) | 108 | 0 | 0 | 36 |
| chr20 | 7399763 | pcsk9 | GGCAGTAAAGTTGCCCCATGTGG | 22/80 (27.5%) | 108 | 0 | 0 | 226 |
| chr20 | 19914504 | pcsk9 | CAACAAGGATCAACATTAGTGA | 14/80 (17.5%) | 40 | 7 | 0 | 29 |
| chr14 | 18273989 | pcsk9 | CCGAATGGGG-AACTTTACTGA | 18/81 (22.2%) | 60 | 2 | 1 | 14 |
| chr24 | 27308362 | pcsk9 | CCGAATGGGG-AACTTTACTGA | 18/81 (22.2%) | 60 | 2 | 1 | 8 |
| chr17 | 26877631 | pcsk9 | GTCAGTAAAGTT-CCCCATTCGG | 19/81 (23.5%) | 65 | 2 | 1 | 6 |
| chr12 | 16562148 | pcsk9 | CCGAATGGGG-AACTTTACTGA | 18/81 (22.2%) | 60 | 2 | 1 | 6 |
| chr20 | 11843066 | pcsk9 | CCGAATGGGG-AACTTTACTGA | 18/81 (22.2%) | 60 | 2 | 1 | 4 |
| chr5 | 9624697 | sh2b3 | CCGCCATGGATTCCAGCGCTCCC | 22/80 (27.5%) | 108 | 0 | 0 | 49 |
| chr15 | 31598137 | sh2b3 | GGGAGCGCAGGAATCCATGGATG | 20/80 (25.0%) | 90 | 2 | 0 | 34 |
| chr10 | 20539546 | sh2b3 | CTAAACACTGGAATCAATGGAGG | 16/80 (20.0%) | 54 | 6 | 0 | 10 |
| chr10 | 15269480 | vldlr | GAGCAGTCTCAGTTCCAGTGTGG | 22/80 (27.5%) | 108 | 0 | 0 | 22 |
| chr5 | 3544150 | ywhaqa | GTGGCCTACAAGAACGTAGTGGG | 19/80 (23.8%) | 81 | 3 | 0 | 167 |
| chr20 | 29566888 | ywhaqa | GTCGCCTACAAGAACGTGGTCGG | 21/80 (26.2%) | 99 | 1 | 0 | 73 |
| chr17 | 32503336 | ywhaqa | GTTGCCTATAAGAACGTGGTGGG | 19/80 (23.8%) | 81 | 3 | 0 | 61 |
| chr6 | 52216272 | ywhaqa | GTGGCCTACAAGAATGTGGTAGG | 19/80 (23.8%) | 81 | 3 | 0 | 28 |
| chr16 | 24684272 | ywhaqa | GTGGCTTACAAGAACGTGGTGGG | 19/80 (23.8%) | 81 | 3 | 0 | 27 |
| chr10 | 17168042 | ywhaqa | CCCACCACATTCTTGTAGGCCAC | 19/80 (23.8%) | 81 | 3 | 0 | 18 |
| chr19 | 12188165 | ywhaqa | CCCACCACATTCTTGTAGGCCAC | 19/80 (23.8%) | 81 | 3 | 0 | 18 |
| chr21 | 34112498 | ywhaqa | CCAACCACGTTCTTGGAGGCCCA | 19/80 (23.8%) | 81 | 3 | 0 | 14 |

**Table S2.** Information about all zebrafish samples examined in this study

| Sample # | Sample ID | Gene targeted | Generation | Age | Fish per sample |
|---|---|---|---|---|---|
| 1 | F01p1_1 | ldlra | F0 | 10 dpf | 30 |
| 2 | F01p1_2 | ldlra | F0 | 10 dpf | 30 |
| 3 | F01p2_1 | nbeal2 | F0 | 10 dpf | 25 |
| 4 | F01p3_1 | sh2b3 | F0 | 10 dpf | 30 |
| 5 | F01p3_2 | sh2b3 | F0 | 10 dpf | 30 |
| 6 | F01p4_1 | ywhaqa | F0 | 10 dpf | 30 |
| 7 | F01p4_2 | ywhaqa | F0 | 10 dpf | 1 |
| 8 | F01s3_1 | sh2b3 | F0 | adult | 1 |
| 9 | F01s3_2 | sh2b3 | F0 | adult | 1 |
| 10 | F01s3_4 | sh2b3 | F0 | adult | 1 |
| 11 | F01s3_5 | sh2b3 | F0 | adult | 1 |
| 12 | F01s3_8 | sh2b3 | F0 | adult | 1 |
| 13 | F01s3_9 | sh2b3 | F0 | adult | 1 |
| 14 | F01s3_11 | sh2b3 | F0 | adult | 1 |
| 15 | F01s3_12 | sh2b3 | F0 | adult | 1 |
| 16 | F01s3_13 | sh2b3 | F0 | adult | 1 |
| 17 | F01s3_15 | sh2b3 | F0 | adult | 1 |
| 18 | F01s3_16 | sh2b3 | F0 | adult | 1 |
| 19 | F01s4_1 | ywhaqa | F0 | adult | 1 |
| 20 | F01s4_2 | ywhaqa | F0 | adult | 1 |
| 21 | F01s4_3 | ywhaqa | F0 | adult | 1 |
| 22 | F01s4_4 | ywhaqa | F0 | adult | 1 |
| 23 | F01s4_5 | ywhaqa | F0 | adult | 1 |
| 24 | F01s4_6 | ywhaqa | F0 | adult | 1 |
| 25 | F01s4_7 | ywhaqa | F0 | adult | 1 |
| 26 | F01s4_8 | ywhaqa | F0 | adult | 1 |
| 27 | F01s4_9 | ywhaqa | F0 | adult | 1 |
| 28 | F01s4_10 | ywhaqa | F0 | adult | 1 |
| 29 | F01s4_11 | ywhaqa | F0 | adult | 1 |
| 30 | F01s4_12 | ywhaqa | F0 | adult | 1 |
| 31 | F01s4_14 | ywhaqa | F0 | adult | 1 |
| 32 | F01s4_15 | ywhaqa | F0 | adult | 1 |
| 33 | F01s4_16 | ywhaqa | F0 | adult | 1 |
| 34 | F03p1_1 | ldlra | F0 | 5 dpf | 30 |
| 35 | F03p1_2 | ldlra | F0 | 5 dpf | 30 |
| 36 | F03p1_3 | ldlra | F0 | 5 dpf | 30 |
| 37 | F03p2_1 | nbeal2 | F0 | 5 dpf | 30 |
| 38 | F03p2_2 | nbeal2 | F0 | 5 dpf | 30 |
| 39 | F03p2_3 | nbeal2 | F0 | 5 dpf | 30 |
| 40 | F03p3_1 | sh2b3 | F0 | 5 dpf | 30 |
| 41 | F03p3_2 | sh2b3 | F0 | 5 dpf | 30 |
| 42 | F03p3_3 | sh2b3 | F0 | 5 dpf | 30 |
| 43 | F03p4_1 | ywhaqa | F0 | 5 dpf | 30 |
| 44 | F03p4_2 | ywhaqa | F0 | 5 dpf | 30 |
| 45 | F03p4_3 | ywhaqa | F0 | 5 dpf | 30 |
| 46 | F12p3_1 | sh2b3 | F1 | 5 dpf | 30 |
| 47 | F12p3_2 | sh2b3 | F1 | 5 dpf | 30 |
| 48 | F12p3_3 | sh2b3 | F1 | 5 dpf | 30 |
| 49 | F12p3_4 | sh2b3 | F1 | 5 dpf | 30 |

| | | | | | |
|---|---|---|---|---|---|
| 50 | F12p3_5 | sh2b3 | F1 | 5 dpf | 30 |
| 51 | F12p3_6 | sh2b3 | F1 | 5 dpf | 30 |
| 52 | F12p3_7 | sh2b3 | F1 | 5 dpf | 30 |
| 53 | F12p4_1 | ywhaqa | F1 | 5 dpf | 30 |
| 54 | F12p4_2 | ywhaqa | F1 | 5 dpf | 30 |
| 55 | F12p4_3 | ywhaqa | F1 | 5 dpf | 30 |
| 56 | F12p4_4 | ywhaqa | F1 | 5 dpf | 30 |
| 57 | F12p4_5 | ywhaqa | F1 | 5 dpf | 30 |
| 58 | F12p4_6 | ywhaqa | F1 | 5 dpf | 30 |
| 59 | F12p4_7 | ywhaqa | F1 | 5 dpf | 30 |
| 60 | F12p4_8 | ywhaqa | F1 | 5 dpf | 30 |
| 61 | F12p4_9 | ywhaqa | F1 | 5 dpf | 30 |
| 62 | F12s3_1 | sh2b3 | F1 | juvenile | 1 |
| 63 | F12s3_2 | sh2b3 | F1 | juvenile | 1 |
| 64 | F12s3_3 | sh2b3 | F1 | juvenile | 1 |
| 65 | F12s3_4 | sh2b3 | F1 | juvenile | 1 |
| 66 | F12s3_5 | sh2b3 | F1 | juvenile | 1 |
| 67 | F12s3_6 | sh2b3 | F1 | juvenile | 1 |
| 68 | F12s3_7 | sh2b3 | F1 | juvenile | 1 |
| 69 | F12s3_8 | sh2b3 | F1 | juvenile | 1 |
| 70 | F12s3_9 | sh2b3 | F1 | juvenile | 1 |
| 71 | F12s3_10 | sh2b3 | F1 | juvenile | 1 |
| 72 | F12s3_11 | sh2b3 | F1 | juvenile | 1 |
| 73 | F12s3_12 | sh2b3 | F1 | juvenile | 1 |
| 74 | F12s3_13 | sh2b3 | F1 | juvenile | 1 |
| 75 | F12s3_14 | sh2b3 | F1 | juvenile | 1 |
| 76 | F12s3_15 | sh2b3 | F1 | juvenile | 1 |
| 77 | F12s3_16 | sh2b3 | F1 | juvenile | 1 |
| 78 | F12s3_17 | sh2b3 | F1 | juvenile | 1 |
| 79 | F12s3_18 | sh2b3 | F1 | juvenile | 1 |
| 80 | F12s3_19 | sh2b3 | F1 | juvenile | 1 |
| 81 | F12s3_20 | sh2b3 | F1 | juvenile | 1 |
| 82 | F12s3_21 | sh2b3 | F1 | juvenile | 1 |
| 83 | F12s3_22 | sh2b3 | F1 | juvenile | 1 |
| 84 | F12s4_1 | ywhaqa | F1 | juvenile | 1 |
| 85 | F12s4_2 | ywhaqa | F1 | juvenile | 1 |
| 86 | F12s4_3 | ywhaqa | F1 | juvenile | 1 |
| 87 | F12s4_4 | ywhaqa | F1 | juvenile | 1 |
| 88 | F12s4_5 | ywhaqa | F1 | juvenile | 1 |
| 89 | F12s4_6 | ywhaqa | F1 | juvenile | 1 |
| 90 | F12s4_7 | ywhaqa | F1 | juvenile | 1 |
| 91 | F12s4_8 | ywhaqa | F1 | juvenile | 1 |
| 92 | F12s4_9 | ywhaqa | F1 | juvenile | 1 |
| 93 | F12s4_10 | ywhaqa | F1 | juvenile | 1 |
| 94 | F12s4_11 | ywhaqa | F1 | juvenile | 1 |
| 95 | F12s4_12 | ywhaqa | F1 | juvenile | 1 |
| 96 | F12s4_13 | ywhaqa | F1 | juvenile | 1 |
| 97 | F12s4_14 | ywhaqa | F1 | juvenile | 1 |
| 98 | F12s4_15 | ywhaqa | F1 | juvenile | 1 |
| 99 | F12s4_16 | ywhaqa | F1 | juvenile | 1 |
| 100 | F12s4_17 | ywhaqa | F1 | juvenile | 1 |
| 101 | F12s4_18 | ywhaqa | F1 | juvenile | 1 |
| 102 | F12s4_19 | ywhaqa | F1 | juvenile | 1 |
| 103 | F12s4_20 | ywhaqa | F1 | juvenile | 1 |

| | | | | | |
|---|---|---|---|---|---|
| 104 | F12s4_21 | ywhaqa | F1 | juvenile | 1 |
| 105 | F12s4_22 | ywhaqa | F1 | juvenile | 1 |
| 106 | F12s4_23 | ywhaqa | F1 | juvenile | 1 |
| 107 | F12s4_24 | ywhaqa | F1 | juvenile | 1 |
| 108 | F01wt_01 | uninjected | F0 | 10 dpf | 30 |
| 109 | wtF0_1-5A | uninjected | F0 | juvenile | 5 |

**Table S3.** Primer information for long-read amplicon re-sequencing of on- and off-target sites

| ID | Forward primer | Reverse Primer | Forward primer | Reverse Primer | Expected size (bp) |
|---|---|---|---|---|---|
| ldlra on-target | GGACTGATCTGCCTCTGAATG | CACTTCCAGCTACCGTGTATG | chr3:19,302,116-19,302,136 | chr3:19,308,406-19,308,426 | 6290 |
| ldlra off-target 1 | TGGAGGTCTGGTTGCTCTAA | GTGTATGGAGAGAGATGGGTTG | chr23:8,186,939-8,186,958 | chr23:8,192,392-8,192,412 | 5453 |
| ldlra off-target 2 | ACGCACACATACCAGTCATC | CACACAGCTAACAACCCTACA | chr16:9,327,571-9,327,590 | chr16:9,333,214-9,333,234 | 5664 |
| ldlra off-target 3 | CCAGAGTAATTGGCAGGTCTAC | GATCGAGCACGCTGTTCTTA | chr10:36,348,775-36,348,796 | chr10:36,354,172-36,354,191 | 5417 |
| ldlra off-target 4 | TCTGCGTGTGAGAGATTGTATG | GCTACCCTTGGTGGTGTAAA | chr19:25,974,714-25,974,735 | chr19:25,980,375-25,980,394 | 5661 |
| ldlra off-target 5 | AACAGACGAGCAGTTGGTATAG | CATAGTAGCCGAGAGGTGTTTC | chr24:7,923,642-7,923,663 | chr24:7,926,264-7,926,285 | 2622 |
| nbeal2 on-target | GAGATGGGTGAATGGAAGGATAA | TCTTGGATGACCTAAGGGTAGA | chr16:16,474,916-16,474,938 | chr16:16,481,105-16,481,126 | 6211 |
| nbeal2 off-target 1 | CGATGGTAGGTGGGTTTCTATC | GGTGAATGGCGCAGTTTATTT | chr12:24,369,712-24,369,733 | chr12:24,375,091-24,375,111 | 5400 |
| nbeal2 off-target 2 | CCTACTCTCTGCCCAAGATAAAG | CGGCTTCAACCACAACATTC | chr8:47,071,177-47,071,199 | chr8:47,076,345-47,076,364 | 5188 |
| nbeal2 off-target 3 | TCAGCATAGCAGCAGCATAG | CTCGGTAGGTCAACACAAGTAG | chr14:16,381,452-16,381,471 | chr14:16,386,500-16,386,521 | 5070 |
| nbeal2 off-target 6 | CCCGCTTGCCTCTTCATATT | CAGAGTGTAGGTTGGCCATTT | chr18:14,201,917-14,201,936 | chr18:14,208,077-14,208,097 | 6181 |
| nbeal2 off-target 7 | CAGAGCCTGAGATGGAAGATATG | GATGTGTCGTACGGGTCAAA | chr10:1,888,407-1,888,429 | chr10:1,893,485-1,893,504 | 5098 |
| nbeal2 off-target 8 | GCTACAAACTCTGTCTCCTCTTC | GCGAGTAATGGTTCCTGTAACT | chr3:35,682,090-35,682,112 | chr3:35,687,058-35,687,079 | 4990 |
| nbeal2 off-target 9 | GTCACAAGCCCTCCACTAAA | GCTCCTCTCACTCAGGATTAAC | chr20:24,395,348-24,395,367 | chr20:24,400,556-24,400,577 | 5230 |
| nbeal2 off-target 10 | GCAACTCTCGAGTCACTTTCT | AGCGGTTATGGTCGCATTAT | chr6:23,358,162-23,358,182 | chr6:23,364,462-23,364,481 | 6320 |
| nbeal2 off-target 11 | GCCTGTGCTCATACGATTCT | CACTGTAAGCCTCTGACACTATT | chr14:11,137,744-11,137,763 | chr14:11,143,443-11,143,465 | 5722 |
| nbeal2 off-target 12 | GCGACCCAGAAGTCTCTTTAC | CCTTCGCTCTCTCTTTCATCTC | chr18:30,357,214-30,357,234 | chr18:30,362,099-30,362,120 | 4907 |
| sh2b3 on-target | GAGATCCGAACTGCTGGATTTA | GTTTGGCTTGCTGAGCTTATG | chr5:9,621,342-9,621,363 | chr5:9,626,092-9,626,112 | 4771 |
| sh2b3 off-target 1 | CTCTTGCTATTTCGGGCTAGTG | GTGTCGTGATGTCTCTGTCTTG | chr15:31,594,333-31,594,354 | chr15:31,601,209-31,601,230 | 6898 |
| sh2b3 off-target 2 | GCAGAACCTCGTGATGTCTATC | CGCTGGTCCACTTCTCAATTA | chr10:20,536,920-20,536,941 | chr10:20,543,191-20,543,211 | 6292 |

| | | | | | |
|---|---|---|---|---|---|
| ywhaqa on-target | CTCTTTCCGCTCCCTATGAATG | GTGATGAGGTGCCAAGCTATAA | chr20:29,564,081-29,564,102 | chr20:29,570,097-29,570,118 | 6038 |
| ywhaqa off-target 1 | TTCACGCTACCTCTGTTCTTG | CGTGCAGCCCTATAGTTCTAAT | chr5:3,540,188-3,540,208 | chr5:3,547,891-3,547,912 | 7703 |
| ywhaqa off-target 2 | CATCGGTGTGCTAAACTGAAATG | GTCTCTCTGCGAACAAGGTAAA | chr17:32,501,047-32,501,069 | chr17:32,506,771-32,506,792 | 5746 |
| ywhaqa off-target 3 | GAAGTGGTACTGCCCTGAATAC | TGCGTCGACACAACCTAATC | chr6:52,212,611-52,212,632 | chr6:52,219,045-52,219,064 | 6454 |
| ywhaqa off-target 4 | CGACACTCAGGTAAGAGAACAC | AGGACCAGAAGATGAAGGTAGA | chr16:24,681,274-24,681,295 | chr16:24,685,847-24,685,867 | 4594 |
| ywhaqa off-target 5 | GACCTCATCATTGGCCCTAAA | GCAGAAGCACTGCAACATAAA | chr10:17,165,822-17,165,842 | chr10:17,170,415-17,170,435 | 4614 |
| ywhaqa off-target 6 | TTACTCGTCCCAACTGCTATTT | GGTCTGGGATACGGTTTGTT | chr19:12,184,966-12,184,987 | chr19:12,190,849-12,190,868 | 5903 |
| ywhaqa off-target 7 | GGCCAAAGATGGAGGATATGAG | GCGTAGAGCGCGATAGTTAAT | chr21:34,109,119-34,109,140 | chr21:34,115,564-34,115,584 | 6466 |

**Table S4.** Distinct CRISPR-Cas9 alleles detected at *sh2b3* on-target, both in F1 individuals and F1 larvae pools, obtained from *sh2b3* founder pair #1.

| Allele | Variant | Del size (bp) | Ins size (bp) |
|---|---|---|---|
| 1 | INSERTION | 0 | 8 |
| 2 | DELINS | 3 | 6 |
| 3 | DELETION | 5 | 0 |
| 4 | DELINS | 6 | 5 |
| 5 | DELETION | 9 | 0 |

**Table S5.** Distinct CRISPR-Cas9 alleles detected at *sh2b3* on-target, both in F1 individuals and F1 larvae pools, obtained from *sh2b3* founder pair #2.

| Allele | Variant | Del size (bp) | Ins size (bp) |
|---|---|---|---|
| 1 | DELETION | 6 | 0 |
| 2 | DELETION | 11 | 0 |
| 3 | DELINS | 1 | 5 |
| 4 | DELINS | 15 | 6 |

**Table S6.** Distinct CRISPR-Cas9 alleles detected at *sh2b3* on-target, both in F1 individuals and F1 larvae pools, obtained from *sh2b3* founder pair #3.

| Allele | Variant | Del size (bp) | Ins size (bp) |
|---|---|---|---|
| 1 | DELETION | 6 | 0 |
| 2 | DELETION | 12 | 0 |
| 3 | DELETION | 10 | 0 |
| 4 | DELINS | 2 | 9 |
| 5 | DELETION | 9 | 0 |

**Table S7.** Distinct CRISPR-Cas9 alleles detected at *ywhaqa* on-target, both in F1 individuals and F1 larvae pools, obtained from *ywhaqa* founder pair #1.

| Allele | Variant | Del size (bp) | Ins size (bp) |
|---|---|---|---|
| 1 | INSERTION | 0 | 2 |
| 2 | DELETION | 21 | 0 |
| 3 | DELETION | 2 | 0 |
| 4 | DELETION | 6 | 0 |

**Table S8.** Distinct CRISPR-Cas9 alleles detected at *ywhaqa* on-target, both in F1 individuals and F1 larvae pools, obtained from *ywhaqa* founder pair #2.

| Allele | Variant | Del size (bp) | Ins size (bp) |
|---|---|---|---|
| 1 | INSERTION | 0 | 4 |
| 2 | DELETION | 3 | 0 |
| 3 | DELINS | 1 | 292 |
| 4 | DELETION | 6 | 0 |
| 5 | DELETION | 13 | 0 |
| 6 | DELINS | 381 | 22 |

**Table S9.** Distinct CRISPR-Cas9 alleles detected at *ywhaqa* on-target, both in F1 individuals and F1 larvae pools, obtained from *ywhaqa* founder pair #4.

| Allele* | Variant | Del size (bp) | Ins size (bp) |
|---|---|---|---|
| 1 | DEL | 6 | 0 |
| 2 | DELINS | 4 | 6 |

*Only one F1 larvae pool was available for this founding pair

**Table S10.** List of off-target events and large SVs in the F1 individuals, which are validated in F1 zebrafish larvae pools obtained from the same mating pair.

| Site | Sample | Variant | Freq | Mating pair | Matched pools | Validated in IGV | Validated in pool |
|---|---|---|---|---|---|---|---|
| sh2b3 on-target | F12s3_7 | 1053 bp del | 0.9728261 | pair 3 | F12p3_3, F12p3_4, F12p3_6 | YES | YES |
| sh2b3 off-target 1 | F12s3_1 | 8 bp del | 0.7214854 | pair 2 | F12p3_7 | YES | YES |
| sh2b3 off-target 1 | F12s3_4 | 8 bp del | 0.7152445 | pair 2 | F12p3_7 | YES | YES |
| sh2b3 off-target 1 | F12s3_5 | 8 bp del | 0.7163842 | pair 2 | F12p3_7 | YES | YES |
| sh2b3 off-target 1 | F12s3_7 | 5 bp ins | 0.5784615 | pair 3 | F12p3_3, F12p3_4, F12p3_6 | YES | YES |
| sh2b3 off-target 1 | F12s3_9 | 8 bp del | 0.7235974 | pair 2 | F12p3_7 | YES | YES |
| sh2b3 off-target 1 | F12s3_10 | 8 bp del | 0.7313567 | pair 2 | F12p3_7 | YES | YES |
| ywhaqa on-target | F12s4_20 | 292 bp ins | 0.8678611 | pair 2 | F12p4_4, F12p4_5, F12p4_6 | YES | YES |
| ywhaqa on-target | F12s4_22 | 292 bp ins | 0.9976162 | pair 2 | F12p4_4, F12p4_5, F12p4_6 | YES | YES |
| ywhaqa on-target | F12s4_23 | 292 bp ins | 0.9773414 | pair 2 | F12p4_4, F12p4_5, F12p4_6 | YES | YES |
| ywhaqa off-target 2 | F12s4_9 | 2 bp del | 0.2772384 | pair 4 | F12p4_9 | YES | YES |
| ywhaqa off-target 2 | F12s4_17 | 3 bp del | 0.8082902 | pair 2 | F12p4_4, F12p4_5, F12p4_6 | YES | YES |
| ywhaqa off-target 2 | F12s4_20 | 3 bp del | 0.6814159 | pair 2 | F12p4_4, F12p4_5, F12p4_6 | YES | YES |
| ywhaqa off-target 2 | F12s4_21 | 3 bp del | 0.8082139 | pair 2 | F12p4_4, F12p4_5, F12p4_6 | YES | YES |
| ywhaqa off-target 2 | F12s4_23 | 3 bp del | 0.8852459 | pair 2 | F12p4_4, F12p4_5, F12p4_6 | YES | YES |
| ywhaqa off-target 2 | F12s4_24 | 3 bp del | 0.4630463 | pair 2 | F12p4_4, F12p4_5, F12p4_6 | YES | YES |

**Table S11.** List of on-target editing events in F1 individuals that are seemingly homozygous and where one single allele is reported in >98% of the sequencing reads.

| Site | Sample name | Mating pair | Variant | Del size | Ins size | Validated in IGV |
|---|---|---|---|---|---|---|
| sh2b3 on-target | F12s3_18 | pair 1 | DELINS | 6 | 5 | YES |
| ywhaqa on-target | F12s4_8 | pair 4 | DELINS | 4 | 6 | YES |
| ywhaqa on-target | F12s4_9 | pair 4 | DELINS | 4 | 6 | YES |
| ywhaqa on-target | F12s4_12 | pair 4 | DELINS | 4 | 6 | YES |
| ywhaqa on-target | F12s4_16 | pair 1 | DELETION | 6 | 0 | YES |
| ywhaqa on-target | F12s4_17 | pair 2 | DELETION | 3 | 0 | YES |
| ywhaqa on-target | F12s4_18 | pair 2 | DELETION | 6 | 0 | YES |
